# Temperature and turbidity interact synergistically to alter anti-predator behaviour in the Trinidadian guppy

**DOI:** 10.1101/2023.04.27.538548

**Authors:** Costanza Zanghi, Milly Munro, Christos C Ioannou

## Abstract

Due to climate change, freshwater habitats are facing increasing temperatures and more extreme weather that disrupts water flow. Together with eutrophication and sedimentation from farming, quarrying and urbanisation, freshwaters are becoming more turbid as well as warmer. Predators and prey need to be able to respond to one another adaptively, yet how changes in temperature and turbidity interact to affect predator-prey behaviour remains unexplored. Using a fully factorial design, we tested the combined effects of increased temperature and turbidity on the behaviour of guppy shoals (*Poecilia reticulata*) in the presence of one of their natural cichlid predators, the blue acara (*Andinoacara pulcher*). Our results demonstrate that the prey and predator were in closest proximity in warmer, turbid water, with an interaction between these stressors showing a greater than additive effect. There was also an interaction between the stressors in the inter-individual distances between the prey, where shoal cohesion increased with temperature in clear water, but decreased when temperature increased in turbid water. The closer proximity to predators and reduction in shoaling in turbid, warmer water may increase the risk of predation for the guppy, suggesting that the combined effects of elevated temperature and turbidity may favour predators rather than prey.

## Background

Environmental conditions can alter sensory abilities and behaviour, with impacts on predator-prey interactions (Abrahams et al., 2007; Domenici et al., 2019, 2007; Rodriguez-Pinto et al., 2020) and hence population dynamics and ecological communities (Dieckmann et al., 1995; Lima, 1998; Orth et al., 1984). For example, in estuaries where water clarity fluctuates widely, visual predators such as fish are often outcompeted by chemosensory predators like crabs, with consequences for the top-down control of food webs (Lunt and Smee, 2020, 2015).

Temperature can influence animals’ metabolism (Clarke and Johnston, 1999), activity levels (Culumber, 2020; Mameri et al., 2020), sensory acuity (Danos and Lauder, 2012; Podolsky, 1994), social interactions (Colchen et al., 2016) and risk-taking behaviours (Biro et al., 2010; Forsatkar et al., 2016). In fish, beside mechanistic changes in motility, temperature fluctuations can have positive or negative effects on anti-predator behaviour depending on a variety of factors, such as predator presence (Weetman et al., 1999), life stage (Colchen et al., 2016), mixed-species shoals (Mitchell et al., 2022; Paijmans et al., 2020) or activity levels of potential shoal mates (Pritchard et al., 2001). This suggests that the effects of increased temperature on behavioural traits are context dependant (Kuruvilla et al., 2022).

Increases in water turbidity can also impact fish behaviour. Turbid water is less clear due to suspended particles that scatter light (Davies-Colley and Smith, 2001). This represents a considerable impairment for species that rely on visual cues, as turbid water can affect reaction distances (Vogel and Beauchamp, 1999), thus the prey’s promptness to escape and how closely a predator can approach its prey before striking an attack (Barrett et al., 1992; Miner and Stein, 1996). Additionally, water turbidity can affect how fish perceive their surroundings. Turbidity can act as cover by concealing prey from predators (Gregory, 1993), which allows prey to spend more time foraging (Wishingrad et al., 2015) in an environment they perceive as low risk. In fact, mortality by fish predators tends to decrease under increased turbidity (Gadomski and Parsley, 2005; Reid et al., 1999). However, turbidity-induced reductions in anti-predator behaviour (Chamberlain and Ioannou, 2019; Snickars et al., 2004) may be maladaptive in some contexts, for example when non-visual predators are considered (Ward and Vaage, 2019) or when turbidity varies alongside other environmental conditions (e.g. habitat complexity (Ajemian et al., 2015)).

Both water temperature and turbidity are increasing in natural habitats. Historic records show that global average surface temperatures have increased over 1°C in the last 170 years and are predicted to increase by a further 3°C by the end of the century (Pörtner et al., 2022). Increasing global temperature is impacting weather patterns such as precipitation strength and frequency, with consequences for river discharge and sediment runoff into waterways (Döll and Zhang, 2010; Mi et al., 2019). Beside climate change, other human-driven processes such as eutrophication and sedimentation are also causing habitats to become progressively more turbid (Xu, 1998). These environmental stressors are increasing globally and simultaneously, often impacting the same aspects of fish behaviour via different pathways. For example, higher activity levels have been observed in fish as a result of both increased temperature (due to higher metabolic rates (Bartolini et al., 2015)) and turbidity (due to an increase in foraging effort (Engström-Öst and Mattila, 2008; Johannesen et al., 2012)). Similarly, shoaling behaviour can increase as a result of enhanced swimming abilities at higher temperatures (Weetman et al., 1998), while it can decrease due to the visual barrier among shoal mates imposed by higher turbidity (Chamberlain and Ioannou, 2019; Kimbell and Morrell, 2015).

However, in a predator-prey context, behavioural responses to the combination of increased temperature and turbidity are unknown. When multiple stressors co-occur, the resulting response may be dominated by the response to only one of the stressors (e.g. comparative effects (Folt et al., 1999)), or by the mathematical sum of the individual stressors’ responses (Piggott et al., 2015). Alternatively, stressors can interact with one another, where the overall response is greater or less than the sum of the two individual responses, creating ecological synergies or antagonisms, respectively (Crain et al., 2008). It is important to carry out controlled experiments with the focus on explicitly evaluating the interaction of multiple stressors, as such responses are otherwise difficult to predict from studies on stressors in isolation (Allibhai et al., 2023; Darling et al., 2010; Ginnaw et al., 2020; Murdoch et al., 2020).

In this study, we tested how the co-occurrence of increased water temperature and turbidity affects fish behaviour in a predator-prey context. Specifically, we assessed the activity levels of both predator (*Andinoacara pulcher*) and prey (*Poecilia reticulata*), the distance between the predator and their prey, and the distance among individuals in the shoal of prey as a measure of group cohesion. In previous studies on guppies, activity (i.e. swimming speed (Kent and Ojanguren, 2015)) and anti-predator behaviours such as shoaling (Weetman et al., 1999, 1998) have been found to increase with increased temperature. For larger fish like South American cichlids such as *A. pulcher*, increases in water temperature have also been linked to higher swimming activity (Brandão et al., 2018). Fish size and water temperature are positively correlated with swimming speed (Kent and Ojanguren, 2015), resulting in prey and, their usually larger, predators being affected by changing temperature to different extents (Ohlberger et al., 2012). Therefore the overall effect of changing temperature may give predator or prey a relative advantage over the other. When subject to water turbidity, guppies have been found to reduce activity levels and shoaling behaviour compared to clear-water conditions (Borner et al., 2015). Similarly, when exposed to higher turbidity, cichlids also reduced overall activity levels, while feeding behaviour increased (Gray et al., 2012, 2011; although see Wing et al., 2021). Based on this earlier work, we predict an increase in temperature and turbidity to have contrasting effects on both predator and prey activity levels and on the distance between individuals in a prey shoal. Due to the impact of these stressors on the same behaviours via different modes of action (Galic et al., 2018), we hypothesise antagonistic interactions for activity levels and shoal cohesion. In contrast, we predict that the increase in temperature and turbidity would have similar effects on the predator-prey distance. This is because with higher turbidity, detection distances will decrease between predator and prey (Sweka and Hartman, 2003), and in warmer water prey will spend more time swimming and engaging in predator inspection behaviours (Weetman et al., 1998). Regardless of the activity levels of the predator, increased activity in prey has been linked to higher encounter rates at high temperatures (Twardochleb et al., 2020). Therefore we hypothesise that the combined response to increased temperature and turbidity may be synergistic.

## Methods

### Study species

Adult male Trinidadian guppies, *Poecilia reticulata* (N=288, mean ± SD = 23.15 mm ± 2.02), were used as prey fish. Guppies were caught in 2019 under license from high predation pools in the Guanapo river in the Trinidadian Northern Range, Trinidad and Tobago, by researchers from the University of Oxford, UK. The guppies were then selectively bred for 3 generations under a controlled breeding plan designed to maintain the genetic diversity of the stock. In 2020, the descendent guppies were transported to the University of Bristol. They were housed in two 90L holding tanks (L x W x H = 70 × 40 × 35 cm) enriched with a sand substrate, artificial foliage and plastic tubes as refuges. The water in the holding tanks was maintained at 23°C, was always kept clear, and the light-dark cycle followed a 12h regime. Fish were fed once a day, after testing, with TetraMin Flakes (Tetra, Melle, Germany).

The blue acara, *Andinoacara pulcher* (N=36, mean ± SD = 11.06 cm ± 1.68), was used as the predator in this study as it is a native predator of the guppy (Deacon et al., 2018). 36 blue acara, bred at the University of Bristol from an original stock provided by the University of Exeter, were divided into groups of 9 in four 45 L holding tanks (L x W x H = 70 × 20 × 35 cm). To allow for the identification of each individual, acaras were given a unique ID code based on their total length and holding tank code. Each holding tank was similarly enriched and maintained as the guppies’ holding tanks. Blue acaras were fed once daily, after testing, with commercially available defrosted bloodworms. This allowed for a minimum non-feeding period of 14 hours before testing.

### Experimental set up

The experimental arena consisted of a transparent cylinder (Ø = 20 cm, H = 15 cm) to house a shoal of 4 guppies, surrounded by an outer ring (Ø = 45 cm, H = 30 cm) where one blue acara was placed, with 10 cm water depth throughout. The predator was netted into the outer ring and left to acclimatise for 10 minutes, after which the prey were netted into the inner circle. The interactions were then video recorded from above (Panasonic HC-VX870, 3840 × 2160 pixel resolution at 30fps) from a 120 cm height for 15 minutes.

The four treatments consisted of a control (water temperature that matched the housing conditions of 23°C and clear water 0 NTU), a warm treatment (29°C, 0 NTU), a turbid treatment (23°C, 15 NTU) and an interaction treatment (29°C, 15 NTU). This range is representative of the conditions encountered in the streams of Trinidad (Magurran, 2005; Magurran and Phillip, 2001) and, although turbidity can fluctuate to levels higher than 15 NTU (Borner et al., 2015), this level of turbidity was selected to allow accurate observation and tracking of the fish’s behaviour. Water temperature was increased using in-water heaters (Hepo HP-608) in a water bath surrounding the test arena. Turbidity was increased by mixing 0.3g of white superfine kaolin clay to the clear water in the experimental arena, assuring the turbidity levels of the outer ring matched the inner circle. The arena was thoroughly cleaned and refilled with clear water after every turbid and interaction treatment test day, and the water was aerated with air stones overnight and in between trials. The temperature was recorded before each trial with an aquarium thermometer. Turbidity levels were recorded at the beginning and end of each test day for control and warm treatments and between each trial for turbid and interaction treatments (2 × 5ml water samples were measured with a turbidity meter, Thermo Scientific Orion AQUAfast AQ3010). Trials were carried out between 8:30 and 17:30 allowing for a minimum of 30 minutes of light before and after testing. The experimental trials were conducted from July to September 2021.

Each test day, all 9 acaras from one of the four groups were tested under one treatment condition, and across the experiment, each acara was exposed to each treatment condition only once in a balanced crossover Latin square design (Figure 1) to minimise order effects. A total of 144 trials were carried out; of these, 9 trials were not included in the final analysis due to technical problems during testing. The 288 guppies were equally divided into two holding tanks (A and B in Figure 1). As individual guppy identities could not be recorded throughout the experiment, the guppy shoals were haphazardly formed for each trial; each guppy was however only tested twice for each treatment. This was ensured by segregating the used guppies after each test day. For example, 36 guppies were haphazardly caught for each test day from holding tank A, until the acara group 1 had been tested in all 4 treatments (144 guppies in total). All guppies from tank A were then tested again with the acara group 3, but with different shoal compositions and following a different treatment order. The same individual fish, for both acaras and guppies, was never tested on two consecutive days, allowing for a minimum of one rest day between testing.

**Figure 1.**
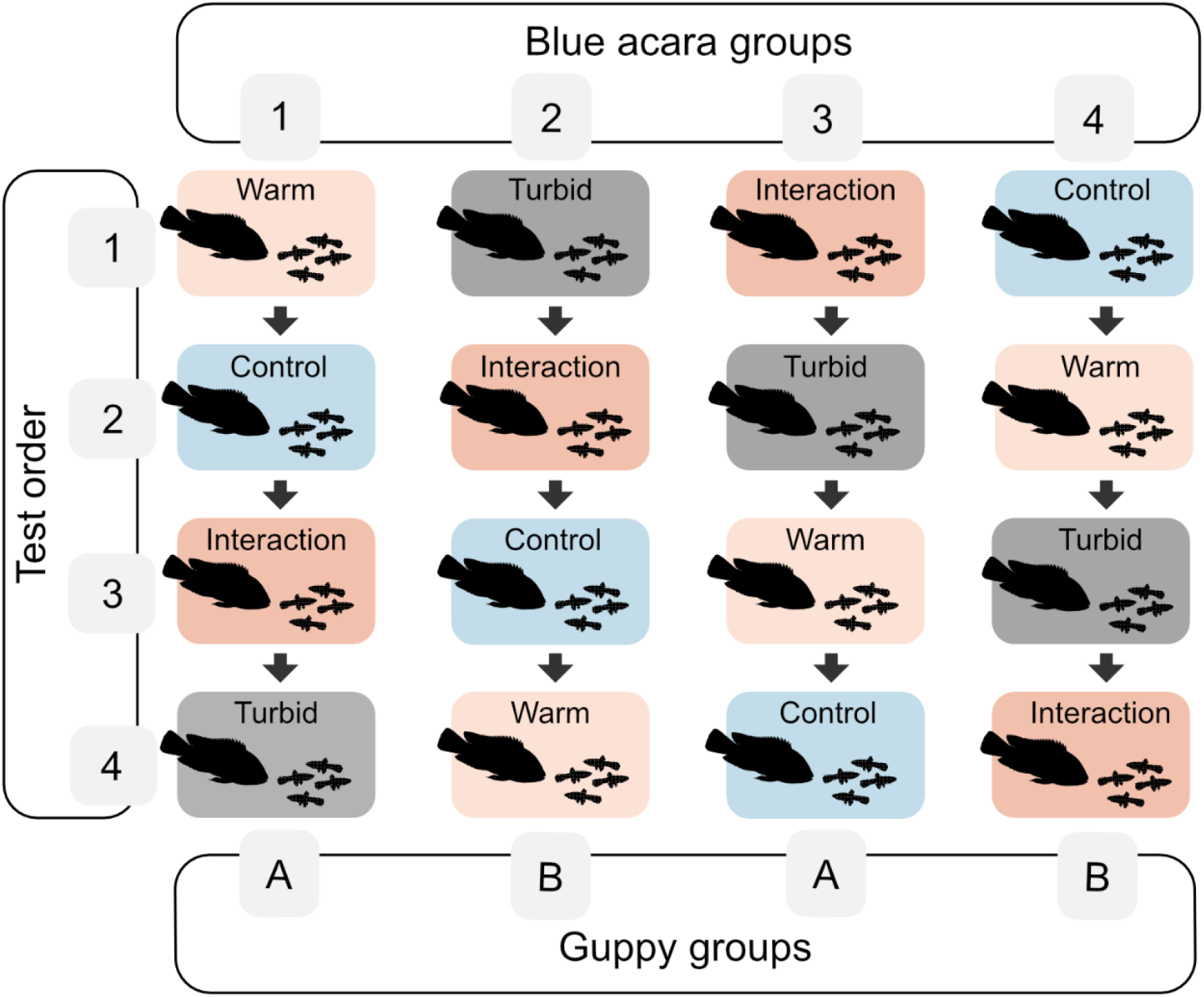
Experimental protocol showing the order in which treatments were presented to each fish group. Predators were divided into groups 1, 2, 3 and 4, while prey fish were divided into groups A and B. The Latin square design allows for each treatment to appear only once in each test order position (1 to 4), therefore providing a unique treatment test order for each holding tank of predator fish.

Both guppies and acaras were housed in clear water, so they were only exposed to turbidity during testing. However, to avoid physiological stress (Brandão et al., 2018; Shah et al., 2017), both species were acclimatised slowly to the increased temperature of the testing condition. The day before each group was due to be tested, fish were moved to temporary holding tanks (200L for 9 acaras, and 54L for 36 guppies). These had separate filtration (EHEIM 2217 for the acaras, Interpret PF1 for the guppies) and heating (Hepo HP-608; 300w for acaras, 50w for guppies) that reached the increased temperature (+6°C) in ∼5 hours, and then allowed for a minimum of 10 hours for the fish to acclimatise to the testing conditions. On the days before a control or turbid treatment, fish were still moved to the temporary holding tanks, but temperature levels were maintained at 23°C.

### Data analysis

The 15-minute videos were converted from .mts to .mp4 using HandBrake (version 1.4.0, https://handbrake.fr/), and then processed with idTracker (Pérez-Escudero et al., 2014) to extract the XY coordinates for each fish in each video frame. The resulting ID-specific tracks were then analysed in R (version 3.6.1) to calculate four behavioural responses: activity levels separately for predator and prey, distance between predator and prey, and inter-individual distance among prey. As a measure of activity, the mean swimming speed was obtained by calculating the distance travelled by each fish in two consecutive video frames, this was then averaged (mean) across the trial. The mean swimming speed was calculated for acaras and guppies separately. The distance between the predator and each prey was calculated and averaged (mean) for each guppy across all the video frames in the trial (i.e. each trial had four mean distances between the predator and prey). As a measure of group cohesion, the distances between each individual guppy and the other 3 guppies were calculated and averaged (mean) across all frames in the trial (mean inter-individual distance).

Each behavioural metric, i.e. the mean prey activity, mean predator activity, mean predator-prey distance and mean inter-individual distance, was included as a response variable in a separate set of generalised linear mixed effect models (GLMMs) using the lme4 R package (Bates et al., 2015). Residuals from all models were checked visually to verify normality and homogeneity of variance. A null model was built for each response variable including only the random effects of the blue acara ID nested in predator holding group. The null model was then compared to seven alternative models each including different explanatory variables: temperature as the only main effect, turbidity as the only main effect, both temperature and turbidity as main effects, an interaction term between temperature and turbidity, minutes from midnight to account for diel variations in fish activity levels, mean guppy total length to account for prey size heterogeneity, and trial order (1 to 4) to account for repeated testing (MacGregor and Ioannou, 2021) (Table 1). The likelihood of each model was then compared using the small-sample correction for the AIC (Akaike information criterion) value. Ranking models with and without explanatory variables of interest allows inference of which variables are more important. If the difference in AIC is greater than 2 units, the model with the smaller AIC value is considered to have strong support as more likely given the data (Burnham et al., 2002). When the null model presents the smallest AIC value, it suggests none of the explanatory variables considered explain variation in the response. To explore possible mechanisms for the trends found from the GLMMs, correlation tests were performed between the mean predator-prey distance and the activity levels of both predator and prey, for these tests the mean predator-prey distance and mean speed for all prey were averaged (mean) per trial so that N=135 for all variables.

**Table 1.**
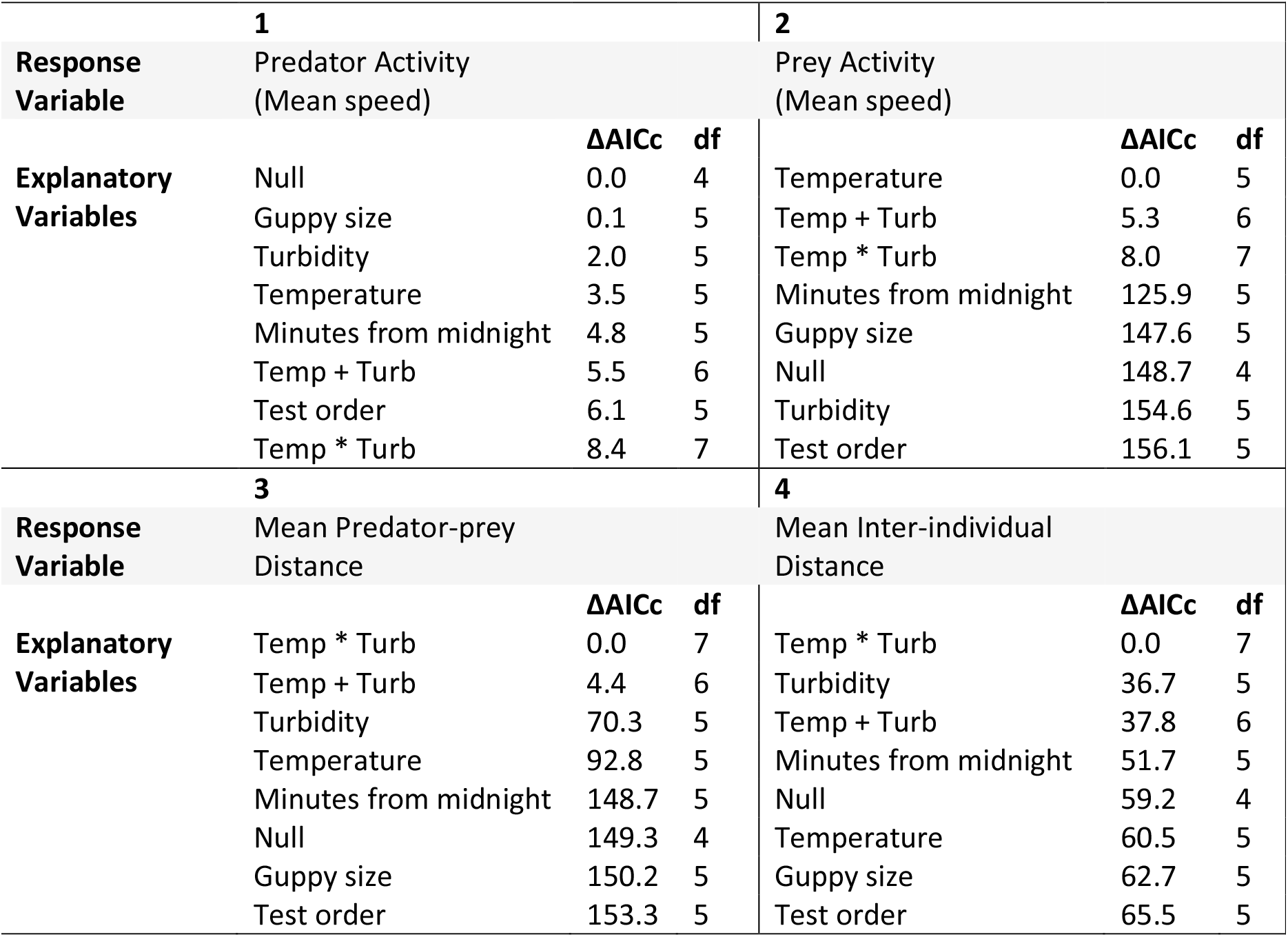
Structure of model comparisons for each behavioural response variable. Each row respresents an individual model varying in their explanatory variables: the plus sign indicates multiple variables are main effects, while the asterisks indicate an interaction term between temperature and turbidity is also included as well as their main effects. Each model also included a nested random effect of 1|Predator group/Predator ID. ΔAICc: Difference in the small sample corrected AIC between each model and the most likely model. df: number of components for each model.

### Ethical note

All methods and procedures were performed in accordance with ASAB/ABS guidelines for the treatment of animals in behavioural research. The research was approved by the University of Bristol Welfare and Ethical Review Body (UIN/21/003). In this research project, it was essential to use live animals to understand the effects of environmental change on predator-prey responses. Care was taken to minimise handling and stress of the study subjects. Turbidity treatments were limited to concentrations of 15 NTU, and temperature treatments to a limit of 29°C. The white kaolin clay used for the turbidity treatments is commonly used for behavioural experiments (Meager and Utne-Palm, 2007; Ohata et al., 2011; Phan et al., 2020). Less than 1g of clay powder per litre of water was used and is representative of ecologically realistic conditions for these species (Magurran, 2005; Magurran and Phillip, 2001). For both species, changes in water temperature were made gradually and within their tolerance range so not to induce physiological stress (Brandão et al., 2018; Shah et al., 2017). After their use in the experiment, the fish were rehoused in the facilities at the University of Bristol for further experiments.

## Results

The model comparisons of the GLMMs supported important interactions between increased temperature and turbidity for the mean predator-prey distance and for the mean inter-individual distance among the guppies; however there was no interaction for the activity response variables (Table 1). The best predictor of variation in the activity response of the prey was temperature as main effect. For the predator however, the null model was the best performing model, supporting a lack of an effect of either temperature or turbidity. The predators’ mean swimming speeds remained similar across treatments (Figure 2a), while an increase in temperature resulted in the prey’s activity levels to increase regardless of turbidity (Figure 2b).

**Figure 2.**
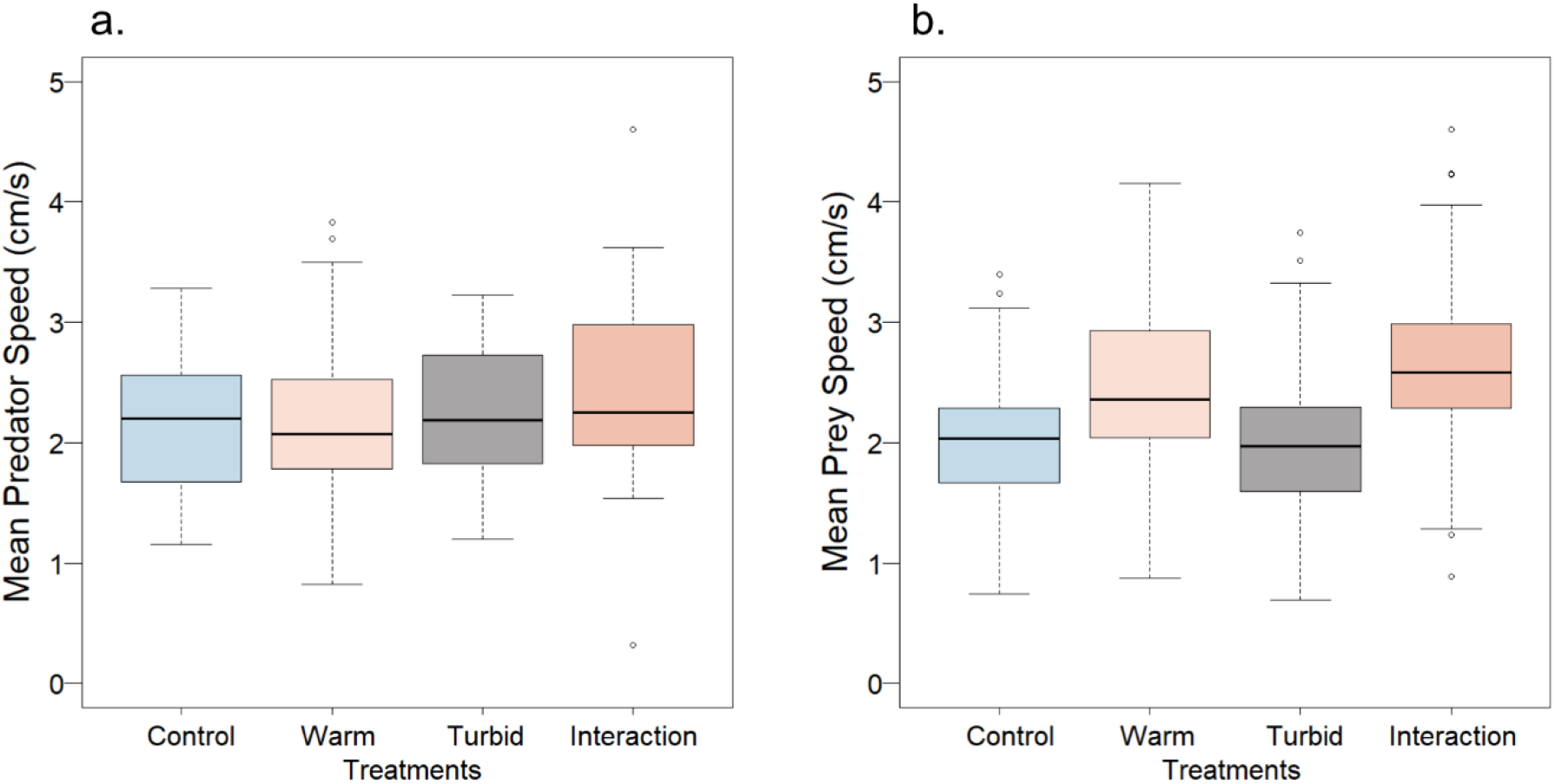
Activity levels for predator (a.) (N=135) and prey (b.) (N=450) across treatment type. Horizontal black lines within the boxes represent the median value. The boxes span the interquartile range. The whiskers extend from the most extreme data point by 1.5 times the interquartile range. The black circles represent outliers.

Compared to the control treatment, the mean distance between predator and prey decreased to a similar extent in both the warm (clear water) and turbid (ambient temperature) treatments. However, it decreased synergistically when turbidity was increased in warmer water, as the decrease in distance was greater than the additive response to the individual stressors (Figure 3a). In contrast, whether temperature increased or decreased shoal cohesion depended on the level of turbidity. In warmer water, the mean inter-individual distance among guppies decreased relative to the control in clear water but increased in turbid water (i.e. in the interaction treatment). At ambient temperature, there was no difference in shoal cohesion in clear and turbid water (Figure 3b). Minutes from midnight had a weak effect only on prey mean speed, all other explanatory variables investigated did not have an effect on the four behavioural responses (Table 1).

**Figure 3.**
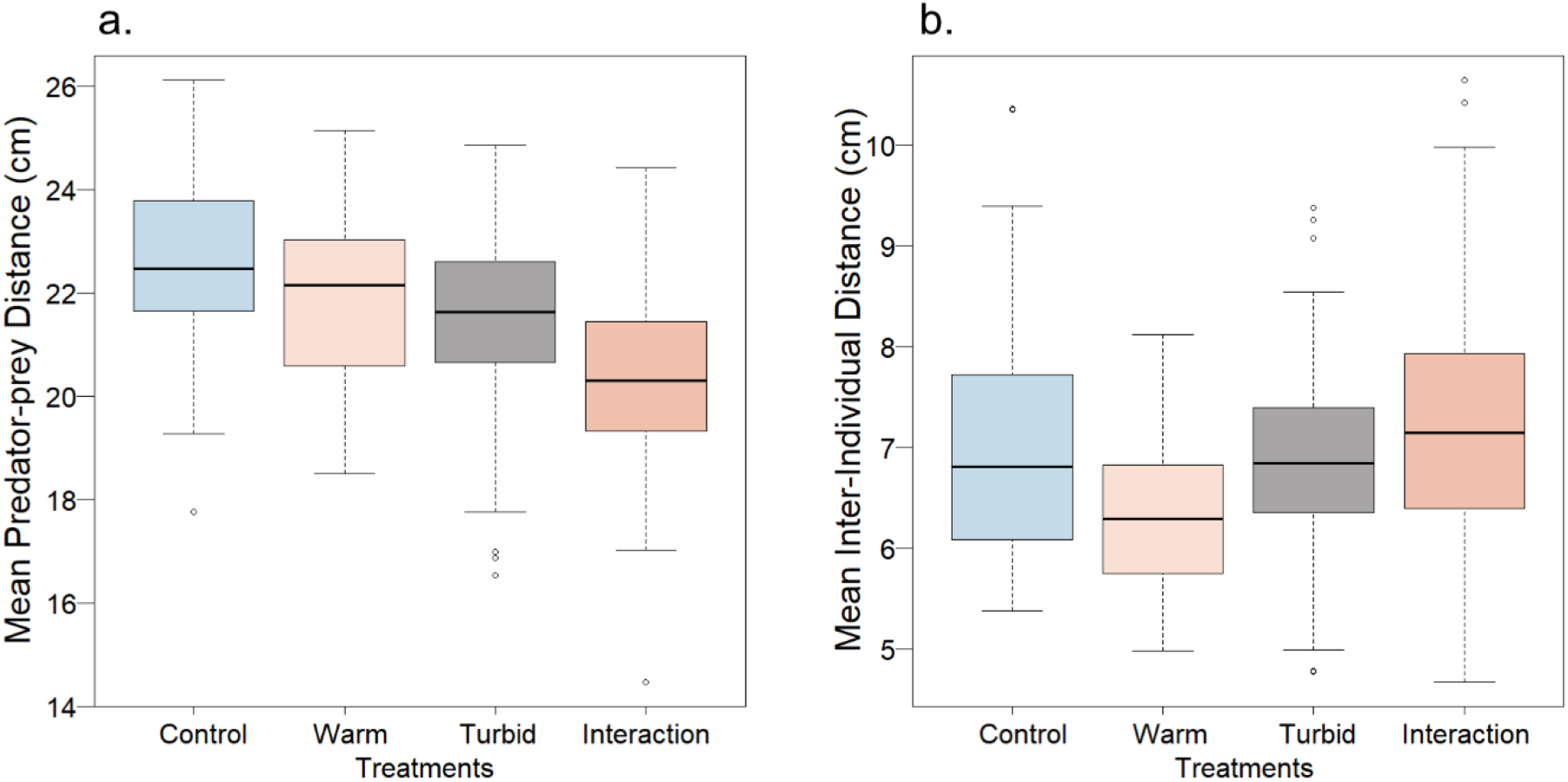
The effect of treatment type on mean predator-prey distance (a.) and mean prey inter-individual distance (b.) (N= 450). Horizontal black lines within the boxes represent the median value. The boxes span the interquartile range. The whiskers extend from the most extreme data point by 1.5 times the interquartile range. The black circles represent outliers.

To provide a mechanistic understanding of these trends, we correlated either predator or prey mean speed with the predator-prey distance for each of the four treatments. Prey mean speed was significantly negatively correlated to predator-prey distance in both the warm (Spearman’s rank correlation, N= 135, r= -0.515, p= 0.003) and interaction treatments (Spearman’s rank correlation, N= 135, r= -0.363, p= 0.030), so that faster prey speed was associated with closer distances to the predator. There were no significant correlations between prey mean speed and predator-prey distance in the control and turbid treatment (Figure 4a). In contrast, a predator’s faster swimming speed was associated with an increased distance between predator and prey in the control (Spearman’s rank correlation, N= 135, r= 0.463, p= 0.008) and warm (Spearman’s rank correlation, N= 135, r= 0.358, p= 0.048) treatments. There were no significant correlations between predator mean speed and distance from prey in the turbid and interaction treatments (Figure 4b).

**Figure 4.**
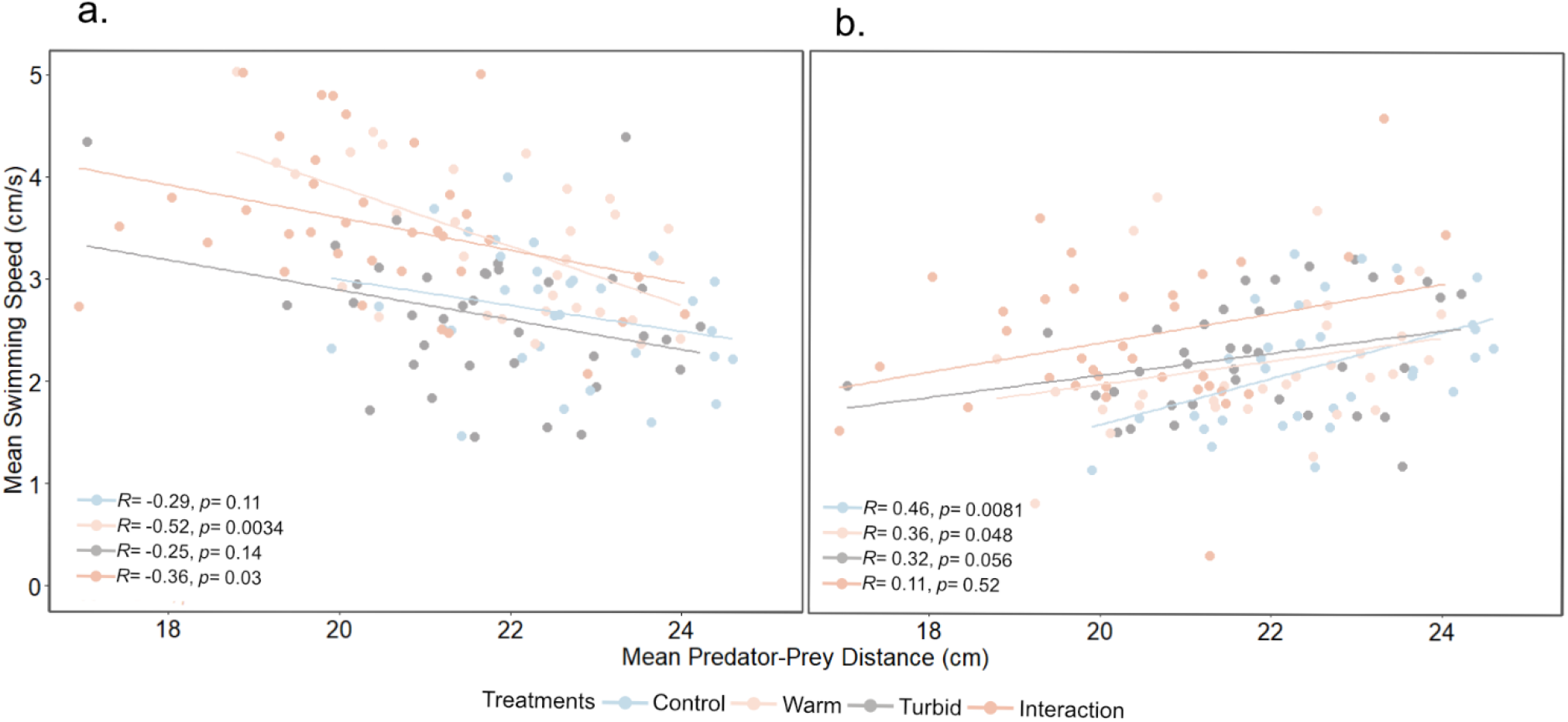
Relationships between mean swimming speed of prey (a.) and predator (b.) and mean predator-prey distance (N= 135). Regression lines, Spearman’s rank correlation coefficient (*R*) and significance of relationship (*p*) are reported for each treatment.

## Discussion

The impacts of multiple anthropogenically-driven environmental stressors on predator-prey interactions have been investigated in a variety of systems (Ajemian et al., 2015; Figueiredo et al., 2015, 2013; Hayden et al., 2015; Horppila et al., 2004; Longmire et al., 2022; McCormick et al., 2018; Nelson et al., 2022). However, the combined effects of increasing water temperature and turbidity on predator-prey interactions are currently unknown, despite the ecological relevance of these stressors and their co-occurrence in our changing world. Here, we found interactions between these focal stressors on the distance between the prey and the predator, and the group cohesion of the prey, while only temperature had an effect on the prey activity levels and neither temperature nor turbidity had an effect on the predator’s activity.

Under control conditions we observed the strongest anti-predator response from the guppies, which on average maintained the furthest distance from the predator compared to the other treatments However, as the activity (swimming speed) of the guppies increased in both the warm and interaction treatments, the predator and prey became closer. This is consistent with previous work demonstrating that increased activity increases encounter rates between predators and prey (Twardochleb et al., 2020). When only turbidity increased, i.e. in the turbid treatment, the predator-prey distance was also reduced, but in this instance activity levels of neither predator nor prey were significantly correlated with the predator-prey distance. The reduced distance compared to control conditions was thus probably due to constrained visibility, reducing the extent to which the guppies could avoid the predator (Miner and Stein, 1996). When increased temperature was coupled with turbidity, we observed a synergistic interaction to further reduce the distance between the predator and prey to a greater extent than the additive effects of the two stressors. This interaction between the effects of greater activity in warmer water and reduced avoidance in turbid water suggests that the prey have further reduced ability to avoid predators in turbid water when they are more active. On the other hand, an increase in the acaras’ swimming speed resulted in greater predator-prey distances, but only in clear water and regardless of temperature. This is likely due to the predator’s greater activity eliciting stronger fleeing responses in the guppies (Cooper, 1998). Additionally, guppies are known to perform predator inspection behaviour (approaching a predator to assess risk and deter attack (Godin and Davis, 1995)) more so when the predator is static rather than actively swimming (Dugatkin and Godin, 1992).

Shoal cohesion increased in warmer, clear water compared to the control, consistent with previous studies where, in warmer water and in the presence of a predator, guppies were found to increase their shoaling behaviour (Li et al., 2022; Weetman et al., 1999). This suggests perception of a higher predation risk when a predator is present in warmer water, possibly because, although escape responses and swimming speed are increased at higher temperatures (Walker et al., 2005), swimming performance is also enhanced in predators (James, 2013). When warmer water was coupled with increased turbidity in our study, however, we observed a mitigating synergism (Piggott et al., 2015) as the direction of the effect of temperature on the inter-individual distance between prey was dependent on turbidity. At higher temperature shoal cohesion increased in clear water compared to control conditions, but shoal cohesion decreased in warmer, turbid water. The increased risk perceived in warm water could have been counteracted by a reduced perceived risk in turbid water, as turbidity can act as cover and reduce anti-predator behaviours (Chiu and Abrahams, 2010; Gregory, 1993; Miner and Stein, 1996). Rather than such an adaptive response, a reduced perception of risk may have been the result of a lack of visual information preventing the guppy to appropriately respond to predator presence, similar to Weetman et al., 1999’s study where guppies shoaled less in warmer water when a predator was absent. Further work is needed to disentangle these mechanisms.

As global temperatures continue to rise, many aquatic ecosystems are expected to experience more frequent and severe temperature fluctuations (Collins et al., 2013; Lee et al., 2019). Similarly, human-driven changes in land use and heightened exploitation of natural resources are leading to increased turbidity of waterways due to sedimentation and run-offs (Davies-Colley and Smith, 2001; Mi et al., 2019). Our study highlights the need for more holistic and integrative approaches to studying the impacts of multiple environmental stressors and their combined effects on interactions between species. Our results suggest that the co-occurrence of increased temperature and turbidity could interact to diminish the anti-predator response of shoaling fish. It is not clear if these changes in behaviour are an adaptation to a more turbid environment which shields prey from predators, or if a lack in risk-awareness would disadvantage prey. Further work that focuses on the predator’s response to prey under these environmental conditions is key to understanding the likely risk to prey under natural conditions (Lima, 2002). Ideally this would be carried out using controlled experiments with freely interacting predators and prey (Romenskyy et al., 2020), including prey mortality, although would have ethical implications. A reduction in anti-predator behaviour is likely to have consequences on prey mortality and hence community dynamics (Wirsing et al., 2021) as prey would become more vulnerable to predator species less reliant on vision and more tolerant of turbid water, such as invertebrates, nocturnal and invasive species (Botham et al., 2006; Chaparro-Herrera et al., 2020; Ehlman et al., 2019; Lunt and Smee, 2014; Sih et al., 2010). Thus, while studies have begun to examine the effects of multiple environmental stressors on predator-prey interactions, even greater ecological complexity may be required by considering prey in a multi-predator species context.

## Acknowledgments

We thank the universities of Exeter and Oxford for the supply of original populations of the study subjects.

